# Direct visualization and tracing of DNA fold in Drosophila embryo

**DOI:** 10.1101/2022.08.16.502515

**Authors:** Fadwa Fatmaoui, Pascal Carrivain, Fatima Taiki, Diana Grewe, Wim Hagen, Burkhard Jakob, Jean-Marc Victor, Amélie Leforestier, Mikhail Eltsov

## Abstract

The chromatin machinery, with DNA-histone assembly and further folding of nucleosome chains, influences DNA availability for functional interactions necessary to the regulation of transcription, DNA replication and repair. Despite models based on *in vitro* studies, the nucleosome chain geometry within the crowded cell nucleus remains elusive. Using cryo-electron tomography of thin vitreous sections, we directly observed the path of nucleosomal and linker DNA *in situ* in intact flash-frozen organism - Drosophila embryo. We quantified linker length and curvature characterizing an irregular zig-zag chromatin folding motif, with a low degree of linker bending. Additionally, nucleosome conformational variability with non-canonical structures and non-octameric particles were observed as individual objects, without structure averaging, highlighting the high structural heterogeneity of native chromatin.

## INTRODUCTION

DNA association with histone proteins organizes eukaryotic genomes into quasiperiodic arrays of nucleosomes connected by a DNA linker (Bilokapic et al., 2018, Olins & Olins, 1978). Nucleosome arrays behave as flexible heteropolymers capable of folding/unfolding upon interactions between the negatively charged DNA and cations and histone tails enriched in positively charged amino-acids. The structural changes of nucleosome arrays influence the local chromatin environment and DNA accessibility and thus play a key role in genome-related processes. However, the precise mechanisms underlying these transitions remain an open question in cell biology (Kornberg & Lorch, 2020). The canonical nucleosome consists of 145 to 147 DNA base pairs wrapped into a 1.7 turn of a left-handed superhelix around an octamer of core histones. Yet nucleosomes are expected pleomorphic, with considerable conformational and chemical variability related to intrinsic dynamics, incorporation of histone variants and post-translational modifications, variations of DNA sequence and binding of additional histone and non-histone proteins (Zhou, et al., 2019, Zlatanova & Victor, 2009). *In vitro*, nucleosome arrays can fold into regular helical superstructures known as 30 nm fibers, of which two families of models exist: solenoids implying linker bending (Robinson et al., 2006) and zig-zags with straight linkers (Schalch et al., 2005), with dependence on nucleosome repeat length (NRL (Routh et al., 2008)), nucleosome structure (Takizawa et al., 2020), and presence/absence of linker histones (Routh et al., 2008). It remains unclear to what extent these models are relevant in the native genome context characterized by variable NRL and DNA sequence, non-uniform binding of linker histones and regulatory proteins, and a highly dynamic nucleosome landscape (Baldi et al, 2020). There is no experimental evidence of 30 nm fibers in native chromatin *in situ*, except in a few highly-specialized cell systems such as rodent rod receptors (Kizilyaprak et al., 2010), granulocytic differentiation (Xu et al., 2021), or fully inactive chromatin of nucleated erythrocytes (Scheffer et al., 2011) and echinoderm spermatozoa (Woodcock, 1994). Instead, recent studies describe chromatin as locally disordered (Maeshima et al., 2019), although high resolution Hi-C experiments and hybrid approaches suggest that local zig-zag and solenoidal folds could exist (Ohno et al., 2019, Risca et al., 2017). Nucleosomes and DNA filaments were imaged *in situ* by transmission electron microscopy (EM) of dehydrated, resin-embedded samples (Kizilyaprak et al., 2010, Ou et al., 2017, Rapkin et al., 2012). But nucleosome conformation and linker DNA geometry depend on electrostatic contacts that need to be preserved, which is achieved by cryo-immobilization of macromolecules in their native hydrous and ionic environment, followed by cryo-electron tomography (Cryo-ET).

A major obstacle in preparing chromatin samples for *in situ* cryo-ET is that conventional plunge freezing methods do not allow vitrification of the thick nucleus in its native state for most higher eukaryotes cell types. This has been circumvented by nucleus or chromosome isolation (Beel et al., 2021, Li et al., 2023), providing *ex situ* information, with modifications of the osmotic and ionic environment. The addition of glycerol has also been used (Chen et al., 2025, Hou et al., 2023), but is known to alter solvation and electrostatics, crucial in biomolecular interactions, especially in the case of chromatin (Bendandi et al., 2020, Diaz et al., 2022, Liang et al., 2001). The vitrification difficulties also limited the choice of the samples accessible for chromatin analysis to “unicellular” systems such as cell cultures, yeasts and Chlamydomonas, with results which might not be fully generalized to differentiated tissues of multicellular organisms. Other limitations are the low signal-to-noise ratio and the anisotropic resolution due to the missing wedge, especially challenging for small and pleomorphic objects such as nucleosomes and DNA filaments. The conventional solution, known as sub-tomogram averaging (STA), has revealed nucleosome structure *in situ* (Cai, 2018, Cai et al., 2018, Chen et al., 2025, Eltsov et al., 2018, Hou et al., 2023a, Wang et al., 2024, Zhou et al., 2024). Classification of nucleosome sub-tomograms suggested the existence of two major open and closed states (Chen et al., 2025, Hou et al., 2023), and the presence of non-canonical structures (Tan et al., 2023). While STA provides information on populations of macromolecular complexes, it lacks precision for analyzing individual structures. This limitation is particularly relevant for DNA linkers, whose path has not been not directly resolved to date. The available intepretations were inferred from the identified angular distribution of nucleosomes (Cai et al., 2018, Hou et al., 2023a). .

In this study, we explored the chromatin landscape in intact multicellular organism, Drosophila embryos vitrified at the late development stage (13-15) when the epigenetic landscapes have been established (Walther et al., 2020). High-pressure freezing (HPF) enabled complete vitrification of embryos without cryo-protectants, keeping chromatin *in situ* in its native state and cellular and organismal context (Bouchet-Marquis & Fakan, 2009, Eltsov et al., 2018). We focused on interphase nuclei of *Drosophila* embryonic brain, and imaged ultrathin cryosections by cryo-ET using contrast enhancement by Volta Phase Plates (VPP)(Danev et al., 2017), well adapted for DNA visualization and nucleosome analysis (Chua et al., 2016), followed by deep learning-based denoising independent on structure averaging (Lehtinen et al., 2018). We were thus able to visualize and analyze the DNA filament and quantify the length and curvature of linker DNA between nucleosome particles. We also detected individually a variety of nucleosome conformations, accommodating different DNA and histone amounts, including non-octameric structures. Taking advantage of the distinctive organization of embryonic *Drosophila* chromosomes, with segregation of constitutive heterochromatin (cHC) into large compact domains, we were able to localize these non-octameric nucleosomal particles in euchromatin and/or facultative heterochromatin nanodomains dispersed in the nucleoplasm.

## RESULTS

### Drosophila embryo as a model system for cryo-ET chromatin analyses

We chose the central nervous system (CNS) of *Drosophila* embryos at late developmental stages (13–15) as a model system because of its many practical advantages. Entire embryos can be vitrified by high pressure freezing (Eltsov et al, 2018, Eltsov et al., 2015), and the CNS can be reproducibly found in the interval 70–100 µm from the embryo’s anterior tip (Supplementary Figure S1AB), thus enabling targeted trimming for cryo-sections (Supplementary Figure S1D). A large relative area is occupied by diploid nuclei that accounts for approximately 50% of the total tissue section area (Figure 1A; Supplementary Figure S1C), thus ideally supporting the targeting of chromatin domains in vitreous sections and facilitating tomographic data collection. Additionally, as shown by freeze-substitution, CNS nuclei reveal the stereotypical organization of *Drosophila* embryo chromatin domains (Figure 1A): compact constitutive heterochromatin (cHC) is attached to the nuclear envelope and associated with the nucleolus (NO), while more dispersed domains of euchromatin (EC) and facultative heterochromatin (fHC) are distributed within the nucleosplasm, letting us distinguish cHC from ECfHC and target these chromatin compartments in cryo-tomograms (Figure 1B, C). This is fully compatible with the fact that the ribosome gene cluster in *Drosophila* is located in pericentric heterochromatin region within the X-chromosome (Hilliker et al., 1980). Domains representing euchromatin and facultative heterochromatin (ECfHC) are small (≤ 300 nm) and located mainly in the central part of nuclei. This chromatin organization is invariably present in CNS nuclei and is also found in the majority of *Drosophila* embryonic tissues.

**Figure 1.**
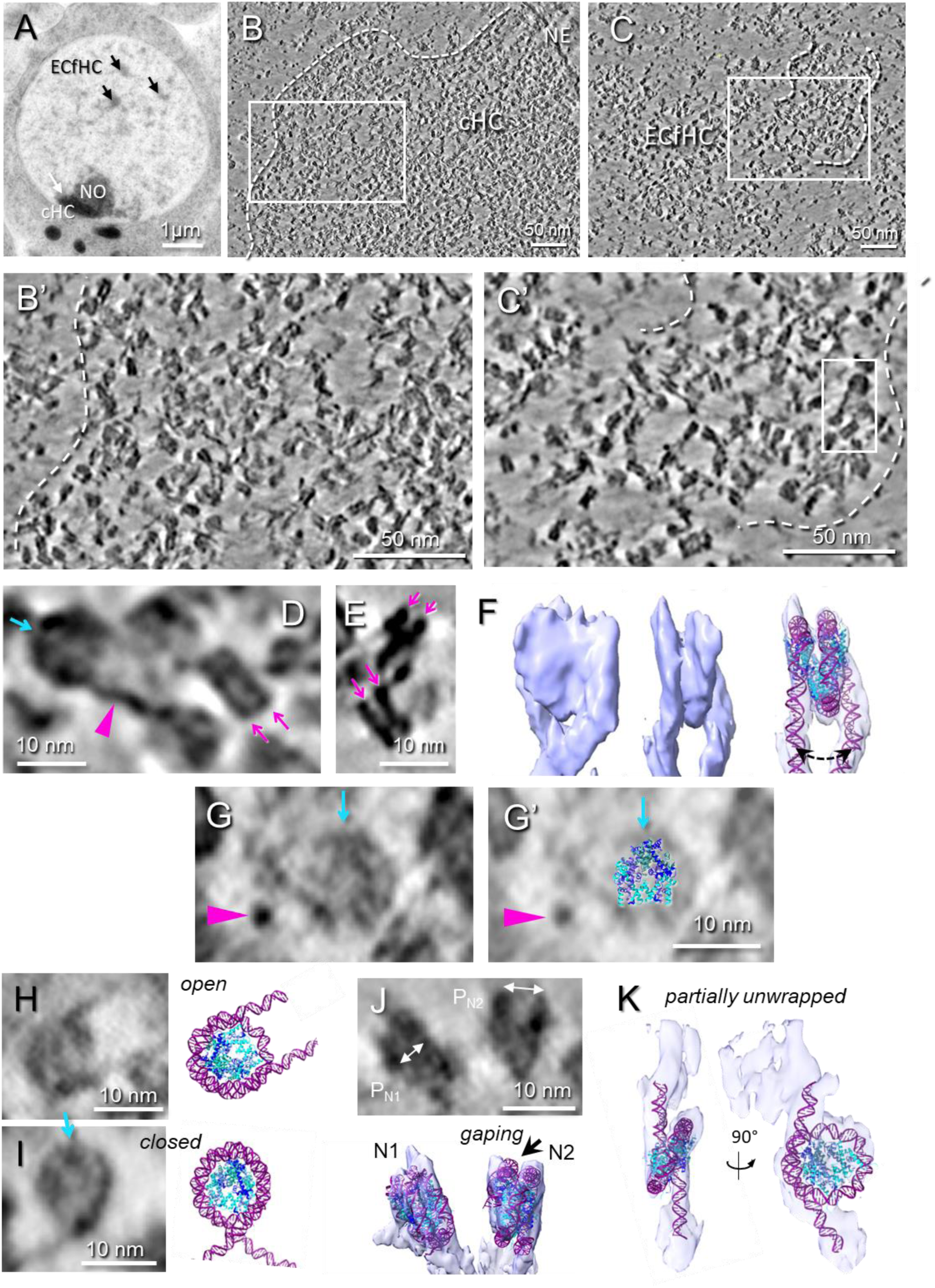
Chromatin visualization enhanced by VPP imaging and computational denoising. (**A**) A section of freeze-substituted embryonic CNS nucleus shows constitutive heterochromatin domain (cHC), nucleolus (NO), and dispersed chromatin domains (arrows) containing ECfHC. (**B, C**) One voxel (4.25 Å) thick tomographic slice of a cryo-tomograms containing cHC (**B**) or ECfHC (**C**) domains after Warp denoising. (**B’, C’**) Enlargements of regions squared in (**B, C**). (**D**) Magnified area from (**C’**) showing nucleosomes in top and side views and linker DNA (magenta arrowhead). The pseudo-dyad axes of the top view nucleosome is indicated by the blue arrow. (**E**) Different side view nucleosomes. The two DNA gyres characteristic of side views are indicated by magenta arrows in (**D, E**). (**F**) Visualization of the complete wrapping of DNA around the histone core in an individual particle, with fitted crystallographic model of the nucleosome (pdb:2PYO). This nucleosome shows a larger lateral spacing between DNA entry and exit sites (dotted double arrow). (**G-I**) Individual top view nucleosomes. In (**G’**) The crystallographic model of the histone fold (from pdb:2PYO) is superimposed. (**G, G’**) A nearby linker DNA seen in cross section is indicated by the magenta arrowhead. A *closed* and an *open* conformation are shown side by side with corresponding models in (**H**) and (**I**) respectively. (**J**) Side views of two neighboring nucleosomes N1 and N2 showing a difference in their intergyral distances (P) are represented (gaping). P_N1_ is very close to that of the canonical nucleosome (∼ 2.7 nm) whereas P_N2_ is larger (∼ 4.0 nm); the cristallographic model (pdb:2PYO) fits poorly in N2 (arrow in isosurface). (**K**) an example of partially unwrapped nucleosome shown as as side and top views, and the corresponding isosurface views with fitted model of hexasome (Bilokapic et al., 2018).

### Visualization of the DNA filament around and between nucleosomes

We recorded VPP cryo-tomograms on either 50 or 75 nm thick vitreous sections (Supplementary Figure S1E). Denoising of tomograms was performed with nonlinear anisotropic diffusion (NAD) filter (Frangakis & Hegerl, 2001) or deep leaning networks based on the Noise2Noise principle (Lehtinen et al., 2018), in Warp (Tegunov & Cramer, 2019) and Topaz (Bepler et al., 2020). In all cases, tomogram reconstructions present a disordered granular aspect typical of chromatin (Figure 1BC). Zooming in reveals nucleosomes (Figure 1B’C’), barely visible in raw reconstructions (Supplementary Figure S2AB, *raw*), but unambiguously identified after denoising (Supplementary Figure S2AB) with their typical contrasted side views dominated by the two DNA gyres drawing characteristic V-, X-, or stripped patterns (Figure 1D, E, J), and circular top views, 11 nm in diameter (Figure 1D). In addition, the density pattern visible in many top view particles is very similar to projection of α-helix histone fold of the octamer (Figure 1G, G’). In the best cases, the complete path of DNA wrapped around the histone octamer may be followed around individual nucleosomes (Figure1F; Supplementary Movie S1).

Importantly, this also reveals DNA linkers connecting nucleosomes (Figure 1D, G, purple arrowheads; Supplementary Figure S2B). The best visibility is obtained by the deep-learning network Warp (Tegunov & Cramer, 2019), but linkers are also revealed by Topaz (Bepler et al., 2020), and nonlinear anisotropic diffusion (NAD) filter (Frangakis & Hegerl, 2001)(Supplementary Figure S2AB), supporting that these linear densities are *bona fide* DNA linkers, not deep-learning artefacts. Although Warp provided the best visibility of these structures, the network training was successful only for reconstructions with <0.6 nm of the mean residual error measured in etomo (Mastronarde & Held, 2017). For volumes with a higher residual error, training produced models that enhanced structural distortions arising from the suboptimal alignment. We found that the best visualization of the DNA filament was obtained on autotrained Warp-denoised tomograms. Topaz training could be successfully performed on all reconstructions, but the structure visibility was not as good as those denoised using Warp (Supplementary Figure S2).

### Conformational variability of individual nucleosomes

We performed STA of manually picked particles from two ECfHC reconstructions (ECfHC1, 549 particles; ECfHC2; 552 particles) and one cHC reconstruction (860 particles), resulting in a 3D nucleosome structure at 13.6 Å resolution in ECfHC1 and 2, and 12.9 Å in cHC (Supplementary Figure S3). Fitting the X-ray atomic model of the *Drosophila* nucleosome core particle (pdb:2PYO) shows that subtomogram averages are very similar to the canonical nucleosome conformation (Supplementary Figure S3). Subtomogram averages show that the structure of nucleosomes is not affected by compression during cryo-sectioning, which confirms previous findings (Cai et al., 2018, Harastani et al., 2021, Harastani et al., 2022).

The analysis of individual nucleosomes reveals rich conformational variability. Disk-like top views of nucleosomes (Figure 1D,G) demonstrate “open” and “closed” conformations (Figure 1H,I). Closed conformations show DNA crossing at the nucleosome entry/exit site (Figure 1I, *closed*). Open conformations show DNA entry and exit points at a distance larger than in canonical X-ray models, revealing partial DNA unwrapping, also called nucleosome breathing (Bilokapic et al., 2018, Buning & van Noort, 2010) (Figure 1H, *open*). This unwrapping could go to a large degree resulting in detaching more than 40 DNA bp (Figure 1K, *unwrapped*) as we can estimate from the fitting of an atomic model. Such unwrapping is characteristic to the hexasome formation via the loss of a histone H2A-H2B dimer demonstrated by single particle cryo-EM *in vitro* (Bilokapic et al., 2018, Zhang et al., 2023), in good agreement with fitting of an atomic model of the hexasome (Figure 1K). Inter-gyre breathing out of the nucleosome plane predicted by simulations (Huertas & Cojocaru, 2021) is also observed at DNA entry and exit sites (Figure 1F, dotted double arrow). Furthermore, some nucleosome side views show inter-gyre distance variation, compatible with gaping (Figure1J, “*gaping”*), i.e. edge-opening, in agreement with FRET experiments *in vitro* (Ngo & Ha, 2015) and our previous findings *in vitro* and *in situ* (Eltsov et al., 2018, Harastani et al., 2021).

Besides nucleosomes, other molecular complexes such as chaperonins and proteasomes are readily recognized in the nucleoplasm (Supplementary Figure S2C).

### DNA linker analysis reveals an irregular zig-zag chromatin folding

Visualization of DNA linkers allows us to trace their trajectories and explore their geometry. Linker paths were traced following the procedure described for cryo tomograms of isolated mitotic chromosomes (Beel et al., 2021) (Supplementary Figure S4). Linkers were identified as fibrillar structures with a diameter of approximately 2 nm in the xy plane, slightly elongated in the z direction due to missing wedge effects. Linkers were observed to be associated with at least one recognizable nucleosome and, under optimal conditions, could be visualized bridging two identifiable nucleosomes, allowing for the reconstruction of the complete linker path (Figure 2A). This class of linkers is hereafter named *2N.* In 46% of the traced linkers (Supplementary Table S1A), only one linked nucleosome can be identified. The linker may end within or close to a density that cannot be assigned to a recognizable nucleosome (Figure 2B; Supplementary Figure S4) and could be another unknown macromolecular complex interacting with chromatin, or correspond to an artefactual density overlap of the tomographic reconstruction (Turonova et al., 2016). The linker may also turn abruptly untraceable (Figure 2B, 1N). We combined all cases where the complete linker length was not defined into class *1N.* We also observed situations where the DNA linker was too short to be traced: successive nucleosomes are then in contact, and DNA can be followed passing from one histone core to the other (Figure 2C, *linker-less*). Altogether, traceable linkers correspond to about 13 % of the amount expected from the number of manually picked nucleosomes (Supplementary Table S1A). Simulated tomograms of synthetic chromatin (Supplementary Figure S5) demonstrate a local variability in the linker path restoration upon reconstruction and denoising, depending on the signal to noise ratio and on chromatin crowding, with partial or even complete loss of the signal in crowded regions (Supplementary Figure S5BC), in good agreement with the situation observed in real data (untraceable linkers or partial linker segments). In addition, to test for a possible bias of the denoising algorithm towards preferential recovery of low-curvature straight linkers versus high curvature linkers, we simulated noisy tomograms containing crowded di-nucleosomes with straight and bent linkers (Supplementary Figure S5, bent linker). We found that both situations have the same signal-to-noise ratio threshold for signal recovery and traceability (0.25; Supplementary Figure S5B). Lastly, we checked that the distribution of the linker “end-to-end” vector directions within the section’s volume shows no statistically significant difference from an isotropic distribution (Supplementary Figure S6), indicating the absence of substantial cutting-induced deformations at the scale of DNA linker.

**Figure 2.**
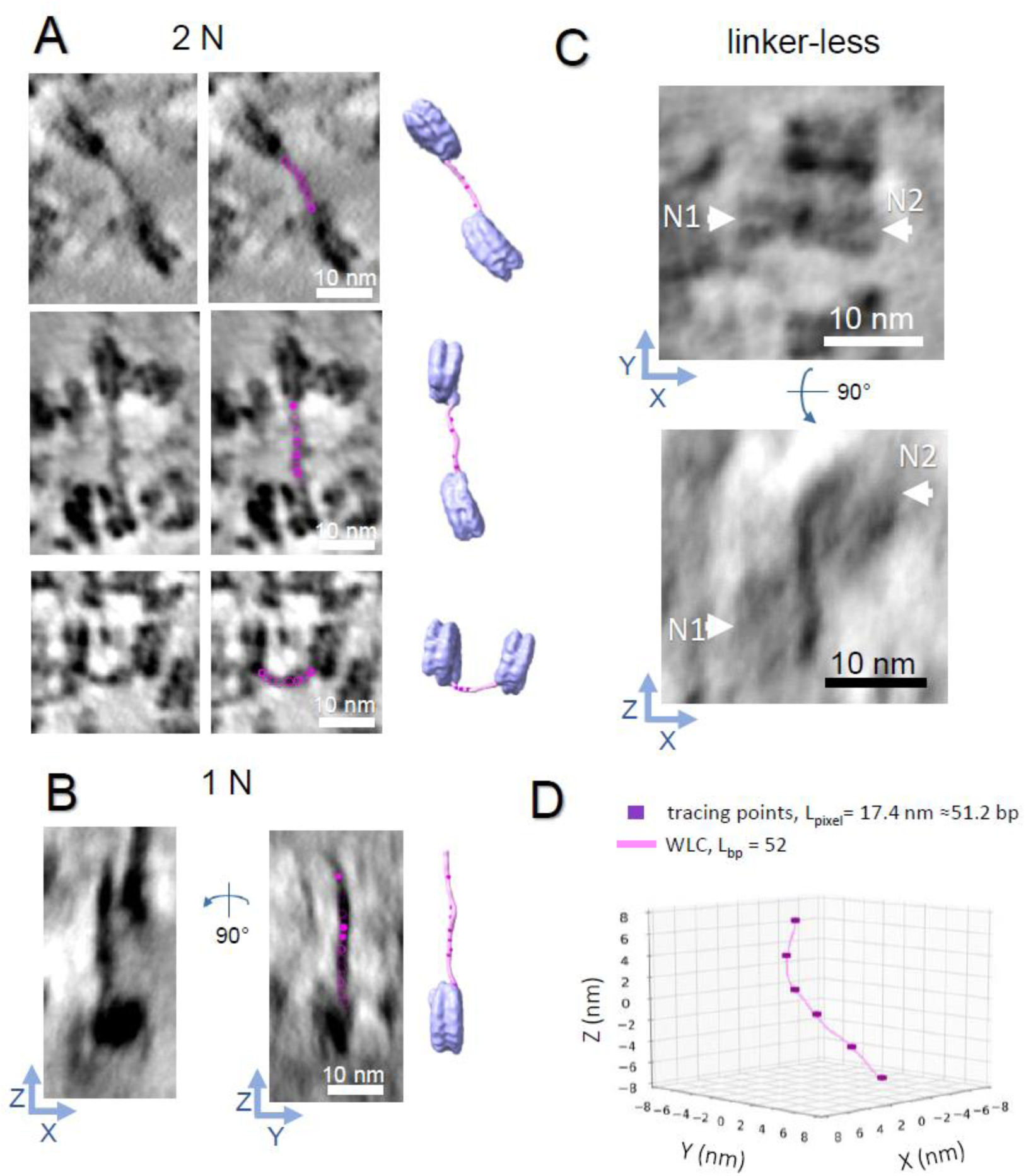
DNA linker tracing and WLC fitting. (**A**) Examples of linker DNA traced between two successive nucleosomes (2N class, **A**) or from one nucleosome only (1N, **B**). Each linker shown first as in one voxel (4.25 Å) tomographic slices (the left column), then with tracing points (purple, the central column) and as a model representing two STA of nucleosomes fitted inside nucleosome densities visible in the given subvolume, and connected with WLCs (pink) fitted to the manual tracing points (purple, the right column). Note that the 1N example shown as two tomographic slices through the central plane of a subvolume is rotated 90° to display a linker that turns abruptly untraceable. (**C**) Example of a linker-less dinucleosome (arrows) shown in side (XY) and top (XZ) views. (**D**) Example of an optimal WLC trajectory (pink) fitted to manually traced points (purple). L_pixel_ represents the length of the polyline connecting tracing points (in nanometers and in base pairs); L_bp_ is the length of the best fitted WLC in base pairs (see Materials and Methods for details).

In order to quantify linker length and curvature, worm-like chain (WLC) models (Doi & Edwards, 1988) with a chain unit of 1 bp and a persistence length of 50 nm were fitted into the traced linkers (Figure 2D). Best fits were processed to calculate linker length and curvature for the *2N* class, curvature only for the *1N* class. Interestingly, the Kolmogorov–Smirnov (KS) test showed no significant difference between linker length and curvature found in ECfHC and cHC (Supplementary Table S1CD). Distributions for the different chromatin compartments are shown in Figure 3A. The mean linker length is 32 bp in ECfHC and 29 bp in cHC (SD 14 bp in both cases, Figure 3A, Supplementary Table S1B), in agreement with previous measurements of 25–40 bp obtained using micrococcal nuclease digestion followed by gel migration and/or sequencing in *Drosophila* cells and larvae (Baldi et al., 2018, Lu et al., 2009). Nevertheless, linkers lengths up to 76 bp and down to a few bp (Figure 3A) are observed, revealing a large local variability. Linker curvature is generally low (mean ∼ 0.10 nm^−1^, SD 0.01 nm^−1^, Figure 3B). No statistically significant differences in linker length and curvature were revealed between ECfHC and cHC (Supplementary Table S1CD).

**Figure 3.**
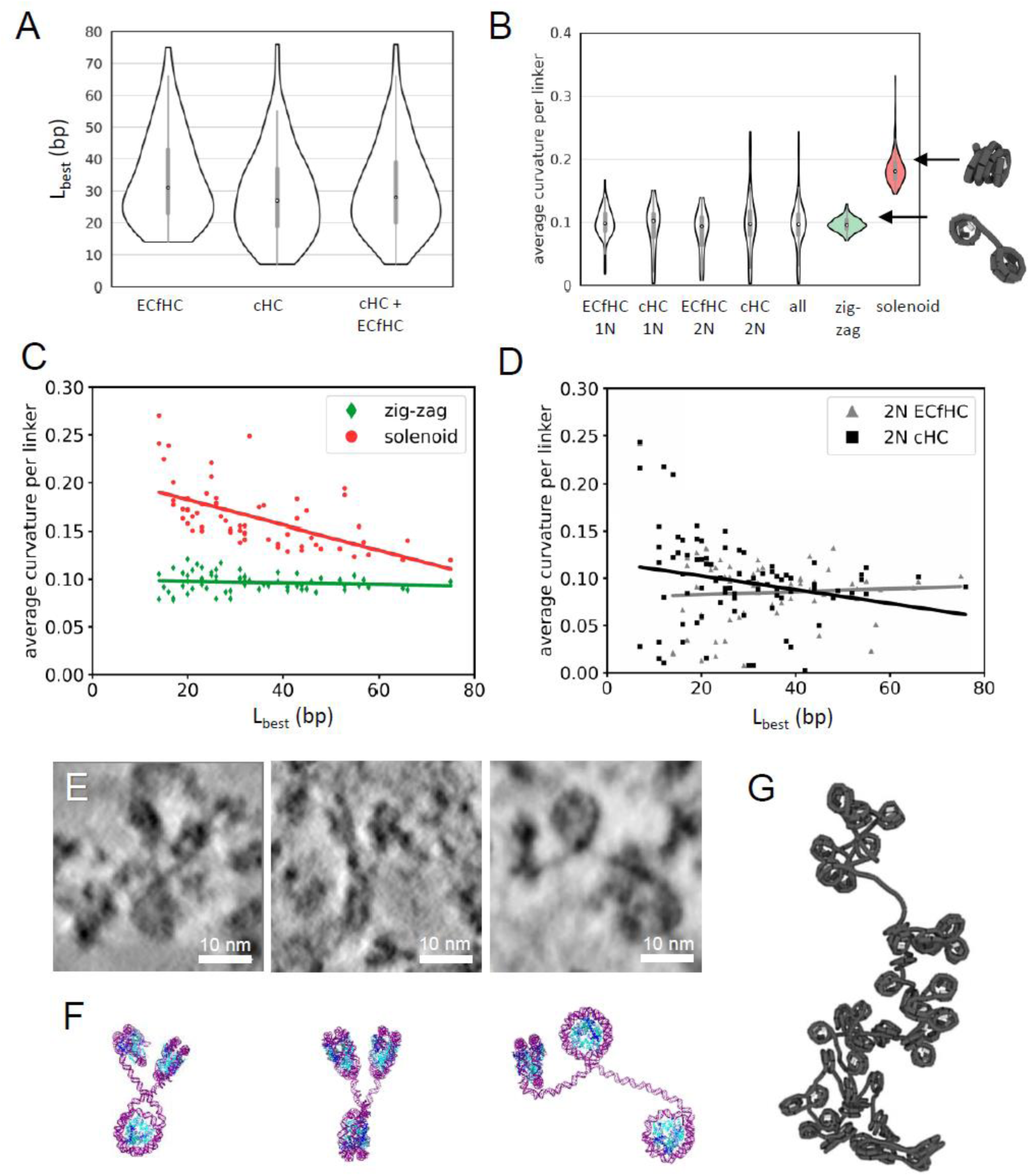
Analysis of DNA linkers demonstrates disordered zig-zag chromatin folding *in situ*. (**A**) Linker length distribution for the 2N class of linkers traced in ECfHC, cHC and all linkers combined. (**B**) Curvature distributions of experimental data plotted separately for 1N and 2N linkers and for all linkers combined (all), presented together with curvature distributions calculated for the zig-zag (green) and the solenoid (red) models computed from the same number of linkers. Schematic views of the minimal structural di-nucleosomes motif of the zig-zag and solenoid models calculated for a linker length of 30 bp (equal to the mean value of experimental data) are shown on the right. (**C, D**) The average curvatures per linker (y-axis) plotted versus its length (x-axis) for the zig-zag and solenoid models (**C**) and the experimental data (**D**). The correlation lines based on the Spearman correlation coefficient show a high significance correlation only for the solenoid model (**D**, *R_SPEARMAN_* = −0.4, p= 7.7 e − 13, see Supplementary Table S2E for details), a low significance correlation for cHC (**C**, *R_SPEARMAN_* = −0.27, p= 0.01), and the absence of correlation between linker curvature and length in ECfHC and the zigzag model (**C, D**). (**E**) Representative 4.25 Å thick tomographic slices illustrating the variety of zig-zag motifs drawn by three successive nucleosomes. (**F**) Molecular models showing our interpretations of views in (**E**). (**G**) Snapshot of a coarse-grained simulation incorporating the measured distributions of linker length and curvature.

To compare our measurements *in situ* with classical zig-zag and solenoid models, we simulated their minimal structural motif as di-nucleosomes, with stacking interaction for solenoids (high linker curvature) and non-interacting nucleosomes for zig-zags (Figure 3B, “zig-zag” and “solenoid”). We plotted experimental linker curvature distributions and compared them to simulated distributions for zig-zag and solenoid fibers. We observed an agreement between the curvature distribution of our experimental data and that of zig-zag models (Figure 3B). In addition to a higher linker curvature, the solenoid model is characterized by a significant negative correlation (−0.40, p = 7.73e-13; Figure 3D, Supplementary Table S1E) between curvature and linker length (*R_SPEARMAN_*) because of a higher degree of bending required for shorter linkers to stack consecutive nucleosomes. Interestingly, while linkers of ECfHC domains, as well as of all linkers taken together show an absence of correlation between linker length and curvature, characteristic of a zig-zag folding, a weak increase of linker curvature with decrease of linker length is indicated in cHC compartment (Figure 3CD, Supplementary Table S1E).

Taken together our results argue against a widespread solenoidal fold and suggest a predominance of zig-zag chromatin geometry, both in ECfHC and cHC, despite indications of increased curvature in short length linkers of cHC. In line with this, wherever two or more consecutive linkers are visualized in the analysed data, they show zig-zag motifs (Figure 3EF).

### Evidence of sub-nucleosomal particles

Besides nucleosomes captured in different conformations and linker DNA, we detected unusual nucleosome-related particles. These structures are connected by DNA linkers to normal nucleosomes and/or each other and visually resemble nucleosomes, but with a key difference: the number of DNA gyre when observed in their side views (Figure 4, Supplementary Figure S7), either one (Figure 4 B-D, Supplementary Figure S7 and S8) or three (Figure 4E, Supplementary Movie S4). They can be unambiguously distinguished from non-nucleosomal bent DNA segments, on account of (i) the presence of an internal cryo-EM density indicating proteinaceous core showing – in some cases - a characteristic histone fold pattern; (ii) their DNA curvature (mean ∼ 0.185 nm^−1^, SD 0.015 nm^−1^, measured on 6 particles), similar to that of the nucleosome (mean ∼ 0.192 nm^−1^, SD 0.017 nm^−1^, measured on 5 particles), and very different from other forms of highly curved DNA such as hairpins (see Supplementary Figure S9). This indicates that 1-gyre particles typify sub-nucleosomes, such as tetrasomes and hemisomes, previously documented *in vitro* and *in silico* (Bancaud et al., 2007, Zlatanova et al., 2009). Both maintain only a single gyre of DNA, while their histone contents are different: the tetrasome contains the complete tetramer of H3/H4 histones, while the hemisome maintains a single copy of each histone. Observed in top view, the tetrasome, with its H3–H4 tetramer, presents a density indentation, along its dyad symmetry axis. This indentation remains discernable on cryo-EM projection patterns simulated at the resolution of our data from an atomic tetrasome model (Figure 4A). The internal density of some of the 1-gyre particles in top views indeed shows such an indentation oriented towards DNA entry/exit place (Figure 4BC, Supplementary Figure S7 and S8) that enables their identification as tetrasomes. In contrast, the modelling of hemisomes predicts their internal density to be asymmetric (Zlatanova & Victor, 2009)(Figure 4A).

**Figure 4.**
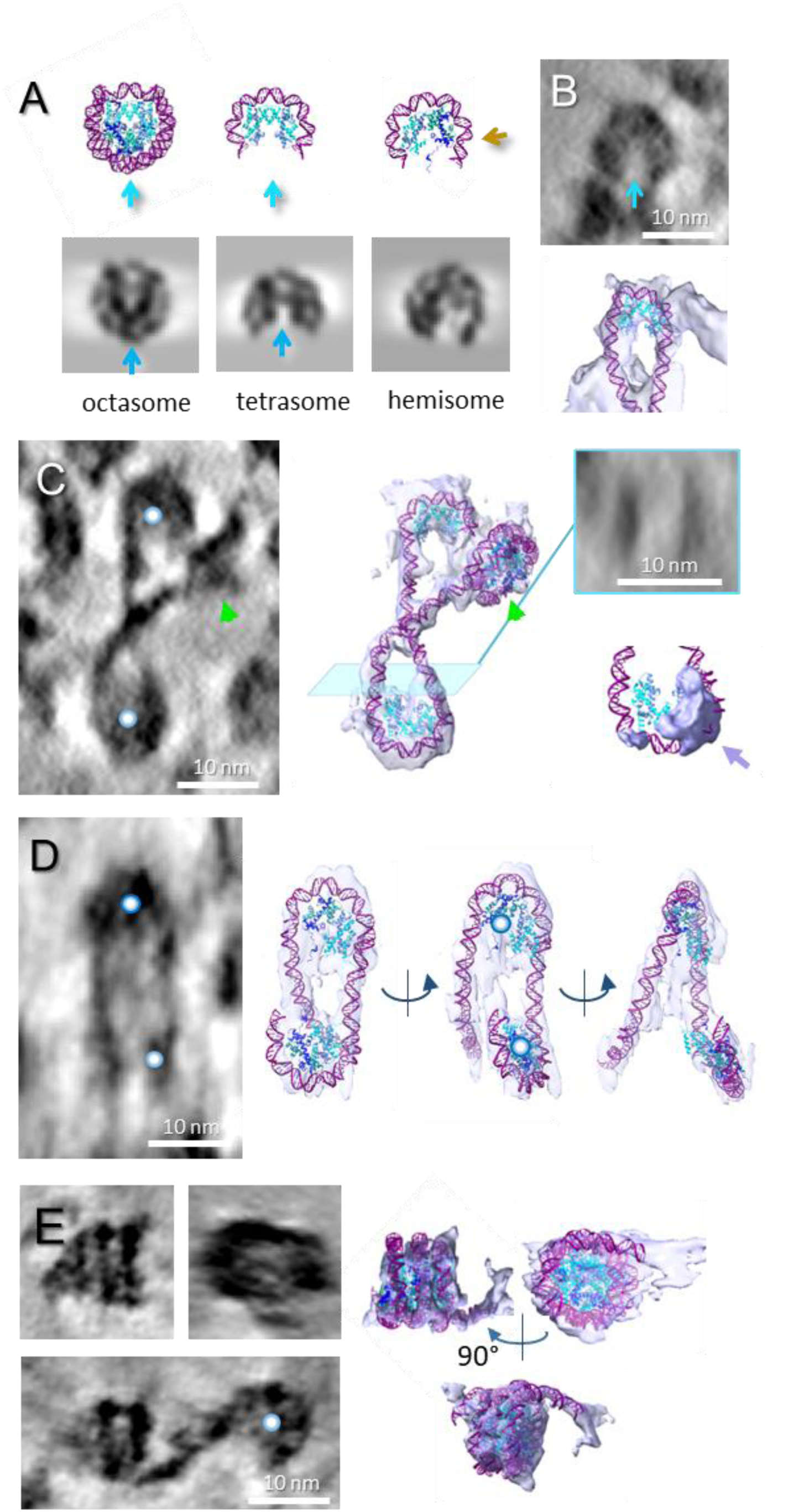
Evidence for unusual nucleosome-related structures in ECfHC domains. (**A**) Atomic models of the complete nucleosome (octasome, pdb: 2PYO) and sub-nucleosomal particles (tetrasome and hemisome). Corresponding simulated cryo-EM densities are shown below. Contrary to hemisome, the H3-H4 tetramer in tetrasome keeps the dyad axis symmetry. Blue arrows indicate the initial dyad axis direction of the nucleosome and tetrasome. In the hemisome, the brown arrow marks the dyad axis orientation of the original octasome before splitting into two halves. (**B**) An example of tetrasome particle showing a symmetry in histone density, shown as a tomographic slice (left) and isosurface with fitted model (right). (**C**) Zig-zag fold formed by two successive tetrasome-like sub-nucleosomal particles (core marked by blue spheres) and a canonical nucleosome (green arrow), shown as a tomographic slice and isosurfaces with fitted models. The transverse tomographic section (inset) corresponding to the blue plane demonstrates that the particle is formed by a single DNA loop. This particle shows an additional density attached to DNA oriented perpendicular to the plane of the DNA loop (blue arrow). (**D**) A pair of hemisomes, corresponding to a split nucleosome with its two halves is represented as a tomographic slice and isosurface with fitted nucleosome and DNA models. The blue spheres indicate the same hemisome in the tomographic slice and models (see also Supplementary Movie S3). (**E**) Three-gyre particle shown as tomographic slices of side and top views and as isosurface views with the fitted model of the overlapping nucleosome (pdb:5GSE). The third modelled DNA gyre does not fully fit the density, indicating conformational difference *in situ* (see also Supplementary Movie S4). Oblique virtual section showing its connection to a sub-nucleosomal (tetrasome-like) particle (blue sphere).

In addition, the histone pattern alone is not the only distinguishing feature between hemisomes and tetrasomes. Hemisomes are predicted to form from nucleosome splitting, as proposed by Zlatanova and colleagues (2009), and are therefore expected to appear in pairs, separated by a short linker (typically less than 50 bp (Zlatanova & Victor, 2009). In contrast, tetrasomes result from the unbinding of 39 bp at both the entry and exit regions of a nucleosome, meaning that two consecutive tetrasomes should be separated by a linker of at least 78 bp. Both scenarios were observed in our data (Figure 4C, D). In Figure 4D (see also Supplementary Movie S3), the two 1-gyre particles not only lack symmetric histone patterns but are also connected by a DNA segment of approximately 25–28 bp—consistent with a pair of hemisomes and incompatible with a pair of tetrasomes. Conversely, in Figure 4C, the histone patterns is more symmetric, and the linker length is around 75 bp — too long for hemisomes but consistent with a pair of tetrasomes.

Interestingly, we also observed a three-gyre or super-nucleosomal particle, which may correspond to an overlapping dinucleosome (Kato et al., 2017), as supported by the fitting of its crystallographic structure (pdb: 5GSE, Figure 4E; Supplementary Movie S4).

All these non-canonical nucleosome-related structures were only found in ECfHC nanodomains, corresponding to about 2% of the nucleosome population. Their small number, combined with their structural diversity, prevents sub-tomogram averaging approaches. A few of them also present additional densities that can be attributed to bound (unknown) proteins (Figure 4C, purple arrow), which, together with their localization in ECfHC domains, suggests that these particles are related to active chromatin regions.

## DISCUSSION

Nucleosome and chromatin structural analysis *in situ* is challenging. With the progresses of cryo-ET, a few recent studies have been able to image chromatin either *in situ* (Cai et al., 2018, Eltsov et al., 2018, Hou et al., 2023a, Tang et al., 2007, Wang et al., 2024, Zhou et al., 2024), of after chromosome or nucleus isolation (Beel et al., 2021, Li et al., 2023). Sub tomogram averaging has revealed a nucleosome average conformation close to the canonical crystallographic one. In all cases, a relatively modest resolution has been reached: 20.0 Å (Harastani et al., 2021); 12.0 Å (Hou et al., 2023a); 12.9 Å (this work), which can be related to conformational variability, as confirmed by dedicated analyses (Harastani et al., 2021, Harastani et al., 2022, Tan et al., 2023)

The use of VPP cryo-ET coupled to deep-learning-based denoising makes possible to visualize the DNA filament in cryo-tomograms, as already shown *ex situ*, by cryo-ET of isolated chromosomes and nuclei (Beel et al., 2021, Hou et al., 2023a). We were thus able to follow DNA trajectories *in situ* within native Drosophila embryo. The DNA path can be visualized, wrapped around and between nucleosome particles, revealing both local chromatin fold and nucleosome conformations at the individual level. The observation of individual nucleosomes not only confirms previous findings of gaping particles (Eltsov et al., 2018, Harastani et al., 2021), but let us to describe various breathing modes (Figure 1FK), and provides direct evidence that nucleosomes accommodate variable DNA length. Between particles, beyond the simple visualization of DNA linkers, we were able to trace the DNA filament. Only 10-15% of the linker paths were determined, due to a combination of crowding (Supplementary Figure S5), presence of a number of proteins bound to DNA and missing wedge effects in tomographic reconstructions. Although this analysis describes a subset of all linkers, a bias toward under- or overestimation of either length or curvature is unlikely because of the detection of very diverse linker lengths and the finding of both strongly curved or nearly straight ones (see Figure 3B). In addition, we were able to visualize directly and trace the zig-zag folding path over 3 to 4 successive nucleosomes.

Altogether, our data reveal a disordered zig-zag folding (Figure 3E-G, Supplementary Movie S2), evocative of the conformations found by Bednar et al. in their seminal observations of oligonucleosome solutions (Bednar et al., 1995). Zig-zag paths were also traced in isolated mitotic chromosomes (Beel et al., 2021) or observed in inactive erythrocyte nuclei (Li et al., 2023), as well as inferred from nucleosome relative positions and orientations at the nuclear envelope periphery of human interphase nuclei (Cai, 2018, Hou et al., 2023). A disordered zig-zag fold, with variable linker length and lack of extensive order is therefore a reasonable model of chromatin folding in interphase nuclei *in situ*. Here, the large variation of DNA linker length (from nearly zero to about 80 base pairs) results in an especially irregular fold, which accounts for the lack of regular fiber coiling motif (Bascom et al., 2017, Collepardo-Guevara & Schlick, 2014, Maeshima et al., 2019). In ECfHC we found no correlation between linker length and curvature, indicating a freely-fluctuating DNA and an absence of geometric constraints, for example stacking of consecutive nucleosomes along the DNA strand like in solenoid type of compaction (Figure 3CD, Supplementary Table S2E). The zi-zag fold minimizes energy penalties for linker bending (Brouwer et al., 2021); it is compatible with the high dynamics of nucleosomes at the local scale and interdigitation of neighboring fibers leading to molten globule-like domains (Maeshima et al., 2019). The low significant correlation between linker length and curvature revealed in cHC data might indicate the accidental presence of stacking interaction between consecutive nucleosomes resulting in linker bending, that might be associated with formation of internucleosome bridges by HP1 (Machida et al., 2018).

In line with this dynamic fluctuating chromatin fold, we observe multiple nucleosome conformations. In particular, closed conformations with DNA crossing at nucleosome entry/exit site coexist with open ones showing variable DNA wrapping/unwrapping. This can occur due to chromatin folding fluctuations and/or the presence of other molecular players, for example bound linker histones resulting in the closed conformation of a chromatosome (Hayes et al., 1994), or chromatin remodeling complexes leading to open reaction intermediates (Dodonova et al., 2020, Farnung et al., 2017, Willhoft et al., 2018). Interestingly, STA obtained by (Cai et al., 2018, Chen et al., 2025, Hou et al., 2023a), have also revealed closed average conformations, consistent with the presence of H1 in a chromatosome.

The *in situ* structural fluctuation of nucleosomes goes beyond the conformational changes of canonical nucleosomes, revealing a distinct set of single-gyre particles. While our resolution does not allow precise identification of their protein components, their DNA curvature and protein density patterns suggest they include tetrasomes and hemisomes (Zlatanova et al., 2009). Additionally, we obtained initial evidences for the presence of triple-gyres particles that might correspond to the overlapping dinucleosome reconstituted *in vitro* (Kato et al., 2017), in which an octasome contacts a hexasome (nucleosome lacking one histone dimer).

Interestingly, all these nucleosome-related structures were found in ECfHC nanodomains. Although we cannot discriminate between EC and fHC, this finding suggests active structural transitions with nucleosome assembly-disassembly and sliding occurring during remodelling and transcription (Bruno et al., 2003, Kobayashi & Kurumizaka, 2019, Kornberg & Lorch, 2020, Petesch & Lis, 2008, Ulyanova & Schnitzler, 2005, Zlatanova & Victor, 2009). Indeed, sub-nucleosomal structures were detected in dynamic chromatin of yeast interphase nucleus (Rhee et al., 2014). Among these, nucleosomes splitted into pairs of hemisomes were recently detected bound by the transcription factor Oct4 (Nocente et al., 2023), and overlapping dinucleosomes may be present downstream of transcription start sites (Kato et al., 2017). The observation of these labile states *in situ* highlights the potential of denoising-enhanced cryo-ET to address functionally-relevant chromatin reorganization directly in the context of cell nucleus.

## MATERIAL AND METHODS

### Reagents and tools

All reagents and commercial instruments are listed in the Reagents and Tools table

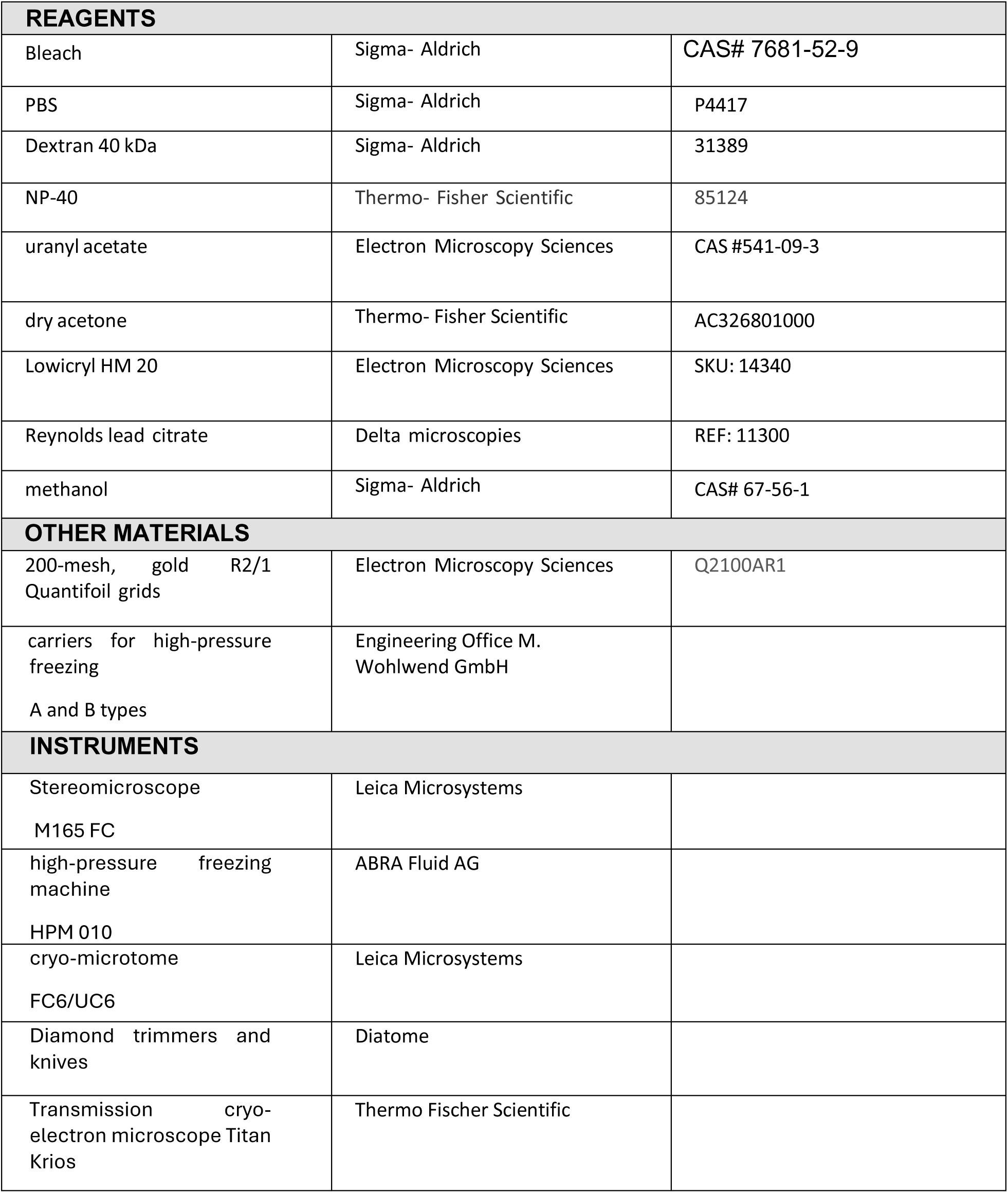

### Biological Resources

*Drosophila melanogaster* (Bloomington Stock number 30564; Dept Biology, Indiana University, Bloomington, USA) were maintained in a standard Bloomington medium.

### Statistical Analyses

Two-sample Kolmogorov–Smirnov(Hodges, 1958) test and Mann-Whitney U Test (Fay & Proschan, 2010) were used. Details and the null hypothesis are described in the corresponding sections of Methods and Methods, and legends of Supplementary Figure S6 and Supplementary Table S1.

### Drosophila embryos preparation and vitrification

Drosophila embryo collection and vitrification were performed as described in Eltsov et al. (Eltsov, 2018, Eltsov et al., 2015). Embryos were collected on apple agar plates, dechorionated in 50% (v:v) bleach (Sigma-Aldrich, St. Louis, USA), washed with PBS and sorted under a stereomicroscope (M165 FC; Leica Microsystems, Wetzlar, Germany). Embryos of developmental stages 14–15 (Campos-Ortega, 1985) were selected and transferred into a drop of the PBS solution containing 25% dextran (40 kDa, Sigma-Aldrich) and 0.25% of NP-40 (Sigma-Aldrich). After < 30 seconds incubation, the embryos were transferred into a 0.1 mm indentation of 3 mm Type A gold-plated copper carriers (Engineering Office M. Wohlwend GmbH). The anterior pole of embryo head was oriented towards the periphery of the carrier (Supplementary Figure S1AD), the carrier was filled with the PBS/Dextran/NP-40 solution and covered by the flat side of Type B carrier (Engineering Office M. Wohlwend GmbH). This sandwich was processed for high-pressure freezing with an HPM 010 machine (ABRA Fluid AG, Widnau, Switzerland). The addition of NP-40 surfactant was necessary for compensation of hydrophobic properties of the embryonic vitelline membrane, thus enabling embryos to be fully surrounded by the Dextran/PBS/NP solution. The wax layer of the vitelline membrane of embryos is not liquid permeable (Rand, Kearney et al., 2010), therefore the Dextran/PBS/NP-40 solution cannot penetrate into the perivitelline liquid and cause any osmotic effect on embryos, whose development occurs normally with larvae hatching.

### Freeze-substitution and conventional TEM analysis

Freeze-substitution was performed as described in Eltsov et al. (Eltsov et al., 2015). The high-pressure frozen embryos were transferred into the chamber of a Leica EM FC6/UC6 cryo-microtome (Leica Microsystems) precooled to –145°C and were punctured with a fine needle (precooled in liquid nitrogen) outside the head area to pierce the vitelline membrane, improving the permeability of the freeze-substitution and embedding media into the embryo tissues. Embryos were then transferred into the AFS-2 apparatus equipped freeze-substitution processor unit (Leica Microsystems) precooled to –90°C. Freeze-substitution began with incubation in glass-distilled acetone (Electron Microscopy Sciences, Hatfield, PA, USA) in the presence of 0.5% uranyl acetate (Electron Microscopy Sciences). After 48 h, the temperature was raised to –45°C at a rate of 2°C/h. After 16h at –45°C, the samples were washed three times with dry acetone and sequentially infiltrated with Lowicryl HM 20 (10%, 25%, 50%, 75%, 4 h for each concentration; Electron Microscopy Sciences). Then three incubations with 100% Lowicryl (10 h each) were carried out, followed by ultraviolet polymerization at –45°C for 48 h, followed by warming to 20°C and an additional 48 h of polymerization.

Embedded embryos were sectioned, perpendicular to the anterior–posterior axis, with diamond knives (ultra 35°; Diatome, Nidau, Switzerland) on a Leica Ultracut E microtome (Leica Microsystems). To explore the general anatomy of an embryo head, serial sections of 200 nm were cut starting from the anterior tip of the embryo and collected on microscopy glass slides, stained with toluidine blue; they were observed under a Zeiss Axiovert light microscope (Carl Zeiss, Oberkochen, Germany) using a x100 objective with oil immersion. After this initial analysis, new serial sections from freeze-substituted embryos were collected on slot grids for electron microscopy (Plano, Wetzlar, Germany) coated with Formwar film (Sigma-Aldrich). Sections were contrasted with 2% uranyl acetate in 70% methanol, followed by Reynolds lead citrate. The sections were examined with Tecnai F30 electron microscope (Thermo Fischer Scientific, Waltham, USA) at magnification ×2300 corresponding to 5.3 nm/pixel and images were acquired with a 4 k × 4 k CCD (charge-coupled device) US4000 camera (Gatan, Pleasanton, USA) controlled by Serial EM software (Mastronarde, 2005). A higher resolution nuclear structure mapping was done using 200 nm thick embedded sections (Figure 1A) acquired at magnification x12000.

### Targeted cryo-trimming and vitreous cryo-sectioning

The carriers with high-pressure frozen embryos were transferred into a chamber of a Leica EM FC6/UC6 cryo-microtome precooled to –145°C and mounted inside a flat sample holder so that the anterior tip of the embryo was oriented toward the knife (Supplementary Figure S1D). The flat-edge portion of trim 45° (Diatome) was used to remove the metal wall of the carrier, then the trimming continued, removing 70 µm of embryo material starting from its anterior tip (Supplementary Figure S1D), to gain access to the CNS. A 90° trimmer (trim 90°, Diatome) was then used to cut the edges off the sides, resulting in a 70 μm × 70 μm × 70 μm cubic block containing the brain tissue (Supplementary Figure S1D). The sections were cut with a nominal cutting feed of 50 or 75 nm using a cryo 25° diamond knife (Diatome) and were collected and attached onto C-flat CF-2/1 or Quantifoil R2/1 grids (Electron Microscopy Sciences) with the Leica EM Crion operating in charge mode.

### Tilt series acquisition, reconstruction, and denoising

For cryo-electron tomography, grids with sections were mounted into Autogrid rings (Thermo Fischer Scientific) and transferred into a Titan Krios (Thermo Fischer Scientific), operated at 300 kV, equipped with a Volta phase plate (VPP), a GATAN GIF Quantum SE post-column energy filter and a K2 Summit direct electron detector (Gatan). The section areas, optimally attached to supporting carbon film, were identified by tilting the grid with 10° steps at ×1400 magnification. The regions of the nuclei overlapping with the C-flat holes were then identified by brief screening at ×2300 (Supplementary Figure S1E), and automated tilt series recording was performed using Serial EM software (Mastronarde & Held, 2017). The dose-symmetric recording scheme (Hagen, Wan et al., 2017) was applied within an angular range from –60° to +60°, with a starting angle of 0° and 2° steps at a nominal magnification of ×64000 (2.12 Å/pixel). The data were collected with a Volta Phase Plate (VPP) in close to focus conditions (Mahamid et al., 2016). To avoid over-focusing parts of images at high tilts, –0.25 μm defocus was applied. The electron dose was set to 2.5 e^−^/Å^2^ for individual tilt images, corresponding to a total dose of 152 e^−^/A^2^ for the complete tilt series.

The denoising of the tomograms was performed with the nonlinear anisotropic diffusion (NAD) (Frangakis & Hegerl, 2001) filter of IMOD (Kremer, Mastronarde et al., 1996) and deep learning networks based on the Noise2Noise principle (Lehtinen et al., 2018) available in the Warp (Tegunov & Cramer, 2019) and Topaz (Bepler et al., 2020) software packages. For training data preparation, dose-fractionation frames of the tilt series were aligned using the IMOD Alignframes tool, resulting in three tilt series: the complete one consisting of tilt images containing all frames for each tilt angle (*complete*); a tilt series consisting of tilt images containing only even frames for each tilt angle (*even*); and a tilt series consisting of tilt images containing only odd frames for each tilt angle (*odd*). Then markerless tilt series alignment was performed on the *complete* series and used to align the *odd* tilt series and *even* tilt series. Tomograms were reconstructed using weighted back projection in the etomo program of IMOD with a pixel size of 4.25 Å (Kremer et al., 1996). A SIRT-like filter, equivalent to 10 iterations, was applied to the reconstructions before applying the NAD filer. Reconstructions used for deep learning network training were done with the standard ramp filter. To provide a consistent input range for the network training, the reconstructions were normalized with EMAN2 (Tang et al., 2007) e2preproc.py to a mean density value of 0 and σ = 1.

Fifteen (15) iterations of the NAD filter, with the K value set to 1, were applied to the complete volumes. Noise2Map network of Warp was trained on the *even* and *odd* reconstructions, starting with the initial model noisenet3dmodel_256 using the following parameters: dont_augment, dont_flatten_spectrum, learningrate_start 1E-05, iterations 40000. The training and denoising was performed on a PC equipped with 2 NVIDIA Quadro RTX8000 GPUs. Topaz denoise3d network was trained on even and odd reconstructions of each tomogram of 4.25 Å/px, without an initial model, using the following parameters: topaz denoise3d --optim adagrad --lr 0.0001 --criteria L2 --crop 96 --batch-size 10 --weight_decay 0 --momentum 0.8 --N-train 1000 --N-test 200 --num- epochs 50 --save-interval 10 --device -2 --num-workers 1. The training and denoising with Topaz was supported by computational resources of a high performance computing cluster of IGBMC.

### Subtomogram averaging of manually-picked nucleosomes and X-ray structure docking

Nucleosomes were manually picked from all reconstructions in IMOD after denoising, then the corresponding 64^3^ voxels (voxel size of 4.25 Å) were extracted from the original non-denoised volumes using AV3 toolbox (Forster & Hegerl, 2007). Subtomogram alignment and averaging were performed with SubTomogramAveraging script (https://github.com/uermel/Artiatomi). The initial reference map was simulated from the crystal structure (pdb:2PYO (Clapier et al., 2008)) with 100 Å resolution using EMAN2. Alignment of subtomograms was performed in 20 iterations. A bandpass filter was applied with a low cutoff frequency of 5 reciprocal-space pixels (1/54.4 Å^−1^), a high cutoff frequency of 12 reciprocal-space pixels (1/22.6 Å^−1^), and a Gaussian edge smoothing with a standard deviation of 3 reciprocal-space pixels. Ten iterations of an unconstrained rotational search (three rotational degrees of freedom) were performed with an angular sampling step of 10°, followed by ten iterations of rotational search constrained to 20° for three rotational degrees of freedom with an angular sampling step of 2°. The Fourier shell correlation (FSC) plot calculated in EMAN2 indicated 13.6 Å at the 0.143 cutoff for all three averages (Supplementary Figure S3). The subtomogram average was denoised with Warp before crystal structure fitting.

### 3D subvolume visualization, atomic model docking, linker DNA modelling

The crystal structure of the *Drosophila* nucleosome pdb:2PYO was fitted automatically into the subtomogram average using the FitinMap tool from UCSF ChimeraX (Pettersen, Goddard et al., 2021); Resource for Biocomputing, Visualization, and Informatics at the University of California, San Francisco, USA). For fitting, we set the simulated map resolution of pdb:2PYO to 14 Å according to the resolution of subtomogram averages assessed by FSC (Supplementary Figure S3). The automated fitting gave cross-correlation coefficients of 0.6197, 0.6015, and 0.6183 for ECfHC1, ECfHC2, and cHC, respectively. The threshold level for the map isosurface was set to 2.86 according to the coverage DNA gyres and histone octamer. The same threshold level was applied to all three averages (Supplementary Figure S3).

To generate the chromatin models shown in Figures 2 and 3, subvolumes of 100^3^ voxels (voxel size of 4.25 Å) containing two or three nucleosomes connected by visible linkers were extracted from denoised reconstructions by boxing tool of EMAN2. The cryo-electron microscopy density map was simulated from the crystal structure (pdb:2PYO) with 18 Å resolution using EMAN2. This map was used for docking into the individual nucleosomes within extracted subvolumes, then the atomic model of nucleosome was fitted inside this map. This two-step docking procedure was used because an automated fitting of the atomic model of nucleosomes into tomogram subvolumes resulted in misorientations.

The accuracy of this docking depends on the orientation of the nucleosome relative to that of the missing wedge. The best results for the translational fitting and orientation were obtained for nucleosomes with the rotational axis perpendicular to *z*-axis of the reconstruction. In this case, the rotational axis of the nucleosome could be defined automatically, then the orientation of the dyad axis was manually tuned to fit the linker path. GraphiteLifeExplore (Life Explorer Initiative, https://www.lifeexplorer.info/) was used to model DNA linkers. Following guidance from Larivière et al. (Lariviere et al., 2017), DNA linkers were placed following the isosurface extracted from the 3D subvolume using UCSF ChimeraX. For simulating crowding effects, the nucleosome-linker models were exported as .pdb files.

To generate the tetrasome model (Figure 4ABC), we started with removing H2A/H2B dimers from the pdb:2PYO structure using UCSF Chimera, resulting in the H3/H4 tetramer and DNA. We then trimmed the DNA so that only the DNA region interacting with the H3/H4 tetramer remained. To generate the hemisome model (Figure 4AD), one of all four histones, and the corresponding gyre of DNA, were removed from pdb:2PYO. Next, the remaining DNA was trimmed, leaving only the part interacting with the remaining histones. The resulting tetrasome- and hemisome-like structures were manually fitted to the corresponding 3D subvolumes (Figure 4BD), then DNA entry/exit regions were modelled following the isosurface extracted from the 3D subvolume using GraphiteLifeExplore. Cryo-EM densities of the nucleosome, tetrasome, and hemisome shown in Figure 4A were simulated in ChimeraX using the molmap command at a resolution of 13 Å and a voxel size of 4.25 Å. The missing wedge was then applied using custom MATLAB scripts based on functions provided by the AV3 toolbox.

### Linker and nucleosomal DNA tracing

Using IMOD routines, DNA linkers were manually traced by three independent observers in 2 volumes containing eu/facultative chromatin domains (ECfHC1 and ECfHC2) and 1 volume of heterochromatin (cHC) denoised by Warp. Linkers were identified as the fibrillar objects found in tomograms, with a diameter of 2 nm, and with one or both ends coinciding with a recognizable nucleosome. To trace a linker, each observer chose a few voxels between which the linker appeared as a straight line (Supplementary Figure S4). The centers of these voxels were then stored as the *tracing points* of the linker. In cases where linkers were identified by more than one observer, the tracings were found to be similar and a single consensus list of tracing points was stored. When only one nucleosome could be recognized, the tracing point numbering started from the nucleosome side. In total, 259 linkers were traced, with 140 *double-ended* linkers (2N) having two nucleosomes identified, and 119 *single-ended* linkers having only one nucleosome (1N) – see Supplementary Table S1A.

The DNA path within round top views of nucleosomes, sub-nucleosomal particles, and sharply bent regions (hairpins) was traced by selecting points within the DNA density of the particle (see Supplementary Figure S9).

### Analysis of linker DNA length and curvature

#### Theoretical background

Linkers were modeled as Worm-Like Chains (WLCs). A WLC is a semi-flexible chain that we describe (as usual) as a curve, *r*(*s*), parametrized by its arc length, *s*, and we denote the tangent vector as 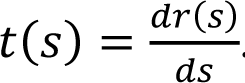. The modulus of the vector 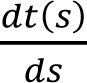 is the curvature κ(*s*) at point *r*(*s*). The elastic energy of the WLC is equal to its bending energy

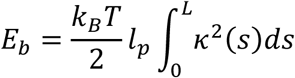

where *l_p_* is the persistence length of the chain, *k_B_* is the Boltzmann constant and *T* = 300*K* is the temperature of the system (before freezing). *L* is the contour length of the chain.

To perform numerical simulations of WLCs, we used a discretized version of the WLC model with segments of 1 bp. In this model, the curvature κ(*s*) is replaced by 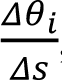, where Τθ_*i*_ is the absolute value of the bending angle between segments *i* and *i* + 1. This discretized version of the curvature requires Τθ_*i*_ ≪ 1 which is fulfilled whenever Τ*s* ≪ *l_p_*. The 1 bp discretization more than satisfies this condition. The bending energy of segment *i* + 1 with respect to segment *i* is:

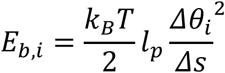

The Boltzmann distribution of the curvature κ(*s*) at any point *r*(*s*) is (following (Rappaport, 2008)):

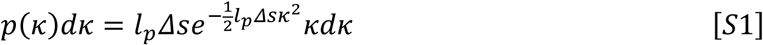

When the linker DNA has an intrinsic curvature, κ_0_(*s*), the above equations need to be generalized. This is the case when the linker is subjected to a given bending torque Γ exerted by its flanking nucleosomes. Then the energy of the corresponding WLC model at mechanical equilibrium is given as a function of the geometric curvature, κ_0_(*s*):

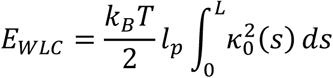

Now, at mechanical equilibrium, the torque Γ is constant along the linker. Since Γ = (*s*), it arises that κ_0_ does not depend on 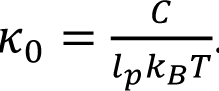. However, thermal fluctuations induce some additional curvature: κ − κ_0_. The resulting Boltzmann distribution of the curvature is then:

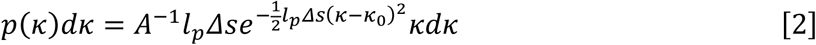

where *A* is a numerical normalization factor:

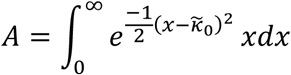

with 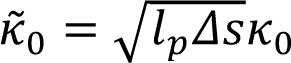.

In this case, the validity of the discretization is subject to the condition: κ_0_Τ*s* ≪ 1, which means that the segment length must be much smaller than the intrinsic radius of curvature of the linker DNA. Again, The 1 bp discretization still satisfies this condition, even in the most stringent case of the solenoid fiber where the intrinsic curvature is κ_0_ = 0,18 *nm*^−1^, i.e. a radius of curvature of ∼5 *nm*.

#### Computing the actual trajectory of the linkers

The trajectory of any linker between two consecutive tracing points is almost straight, hence the sum of the distances between consecutive tracing points is a proxy of the actual contour length of the linker, either single- or double-ended. This sum, expressed in nanometers, is denoted *L*_*nm*_. Then the integer part of the ratio 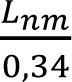, here denoted 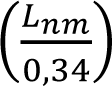, is also a proxy of (and always smaller than) the actual number of base pairs in the linker DNA.

To find the actual trajectory of each linker, we designed the following algorithm to find the WLC conformation(s) that cross(es) all its tracing pixels:

1. We start from a contour length 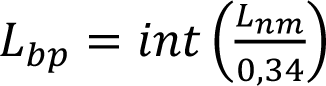
2. We run a WLC simulation with contour length *L*_*bp*_, persistence length *l_p_* = 50 *nm* and 1 bp discretization, and keep the optimal conformations, i.e., those that cross the largest number of tracing pixels.
3. We define the overlap σ as the fraction of tracing pixels that are crossed by the WLC (Γ = 1 when the WLC crosses all tracing pixels). If the overlap Γ < 1, we increment *L*_*bp*_by 1 and we go back to step 2.
4. If the overlap Γ = 1, the algorithm stops and the corresponding value of *L*_*bp*_ is denoted *L*_*bp*,*best*_. This value *L*_*bp,best*_ is the best estimate of the actual number of base pairs in the linker DNA.

Simulations were run for a given number of Monte-Carlo steps MCS = 10^8^ before stopping. For each of the three tomograms (ECfHC1, ECfHC2 and cHC), we ran the above algorithm for 1N and 2N linkers.

#### Computing the contour length of the linkers

We computed the distribution of *L*_*bp*,*best*_ for 2N linkers in each of the three tomograms (ECfHC1, ECfHC2 and cHC) and we characterized each corresponding distribution with its two first moments (mean and standard deviation), as well as skewness and Fischer’s definition of the kurtosis (Supplementary Table S2B). All distributions are clearly not normal. Skewness shows that the *L*_*bp*,*best*_ distributions are strongly asymmetric.

We then investigated whether these three histograms were significantly different or whether they could be put together into a common whole. We therefore applied a Kolmogorov–Smirnov (KS) two-sample test (Hodges, 1958) between any two of these three histograms to test the null hypothesis *H*_0_: “two experimental distributions are actually drawn from the same distribution”. We used the <u>Scipy function</u> (Jones et al., 2001) to run the KS test and compute the corresponding *p*-values. The results of these tests are given in Supplementary Table S2C. The null hypothesis *H*_0_ “the two experimental distributions are drawn from the same distribution” cannot be rejected at the level of significance α = 0.1 neither for eu-nor heterochromatin histograms, therefore the *L*_*bp*,*best*_distributions can be merged for analysis (see Figure 3A).

#### Computing the curvature of the linker and nucleosomal DNA

The algorithm we described above gives access to the whole conformation of the WLC which has perfect overlap Γ = 1 with a given linker. For this best-simulated WLC we computed the local curvature (Τθ_*i*_)⁄Τ*s*, where Τθ_*i*_ is the absolute value of the bending angle between segments *i* and *i* + 1. Curvature is expressed in nm^−1^. We could then compute the average curvature of the linker as the mean of the local curvature along the best simulated WLC. The local curvature is the curvature between two consecutive base pairs. For example, a linker of length 10 bp has 10 – 1 local curvatures. The average curvature per linker is then the average over these 10 – 1 local curvatures. The same approach was used to estimate the average DNA curvature within nucleosomes, sub-nucleosomal particles, and a sharply-bent region (hairpin) (Supplementary Figure S9C). To compare the curvature distributions between nucleosome and sub-nucleosoomal particles, we used Mann-Whitney U test (Fay & Proschan, 2010).

To compute the linker curvature, we considered both 2N and 1N linkers from ECfHC1, ECfHC2, and cHC tomograms because curvature is a local feature, unlike the linker length which is a global feature. We then ran a Kolmogorov–Smirnov two-samples test between any two of the six distributions of the average curvature per linker to test the null hypothesis *H*_0_: “are the values drawn from the same distribution?”. The results of these tests are given in Supplementary Table S1D. The *p*-values showed that we could not reject the null hypothesis *H*_0_. Consequently, we were allowed to combine the curvature distributions for further analysis (see Figure 3B).

#### Coarse-grained model of zig-zag and solenoid fibers

To compare these experimental histograms with the classic zig-zag and solenoid models, we also computed the distribution of the average curvature per linker for these two models (see Figure 3B). For each optimal linker length *L*_*bp*,*best*_that was extracted from one of the three tomograms ECfHC1, ECfHC2 and cHC, we built both a zig-zag and a solenoid fiber, according to the following procedure:

i. A coarse-grained model of a rigid nucleosome is first built at the scale of 10.5 bp. A frame is attached to each monomer of 10.5 bp (corresponding to a length *l* = 3.57*nm* and a diameter *d* = 2*nm*). The tangent follows the centerline of the DNA while the normal vector points towards the minor groove. A linker (with contour length *L*_*bp*,*best*_) exits the first nucleosome and connects to the second nucleosome. The orientation of the second nucleosome is given by the DNA (minor groove) rotational phase. The histone octamer is made of four capped-cylinders to avoid any nucleosome– nucleosome overlap. The system is coupled to a Langevin thermostat at the room temperature *T* = 300*K* (Carrivain, Barbi et al., 2014). Bending torques (the persistence length we used is *l_p_* = 50*nm*) and twisting torques (with persistence length *l*_*t*_ = 85*nm*) mimic the mechanical properties of DNA. Rigid bond constraints enforce DNA connectivity.
ii. We add stacking interaction between two consecutive nucleosome core particles to model a solenoid fiber (one start chromatin fiber, see Figure 3B, *solenoid*). Although there is no stacking interaction for the disordered zig-zag model (see Figure 3B, *zig-zag*), there is still excluded volume between nucleosomes. To minimize end effects, we extract one linker in the middle of the chromatin fiber to measure the average curvature per linker.

Of note, the local curvature density of the linkers in the zig-zag (resp. solenoid) model is in good agreement with the theoretical expression Equation [1] (resp. Equation [2]), as shown in Supplementary Figure S10. Therefore, we are confident that our simulations properly reproduced the curvature behavior of linker DNA in general.

#### Calculation of correlation between linker length and average curvature

Spearman correlation coefficients (*R_SPEARMAN_*) between length *L*_*bp*,*best*_ distributions of 2N linkers from ECfHC1, ECfHC2, and cHC, and their average curvature distributions were calculated using Scipy (seehttps://docs.scipy.org/doc/scipy/reference/generated/scipy.stats.spearmanr.html for more details).

### Simulation of crowding effects on DNA linker visibility after denoising

Tomographic volumes containing chromatin with bent and straight linkers were simulated with custom Matlab scripts based on functions provided by the AV3 toolbox. Initially, atomic models containing two nucleosomes connected with a linker were built in GraphiteLifeExplore and saved as .pdb files. The model of the real volume two-nucleosome connected by the 50 bp linker shown in Figure 2A (upper row) was used for simulation for the case of a straight linker. To simulate the bent linker situation, two nucleosomes were aligned along rotational axes and connected with 60 bp linker DNA (Supplementary Figure S5A). The intrinsic curvature κ_0_ of the straight and the bent linkers were 0 nm^-1^ and 0.13 nm^-1^ respectively. The density map was generated in UCSF ChimeraX using the molmap command with a resolution of 4.25 Å/px. The map was used without applied defocus, and placed into 1024^3^ voxels (435.2^3^ (nm)^3^) volumes, whilst being oriented randomly in all three rotational degrees of freedom. Each volume was filled separately with di-nucleosome densities with either straight or bent linkers. We could achieve the density of 4.2 × 10^5^ nucleosome/(µm)^3^ corresponding to 0.69 mM both for low- and high-curvature linker di-nucleosomes. This value is of the same order of magnitude as the density of compact chromatin *in vivo* measured by fluorescence-based techniques (0.25–0.5 mM) (3,4) To explore the visibility of linkers and nucleosomes at low crowding conditions we generated a volume filled with the same randomly-oriented di-nucleosome densities with the concentration of 5.5 × 10^3^ nucleosomes/µl (9 µM).

To mimic the sectioning process, we extracted the central slice of the simulated volumes measuring 435.2 × 435.2 × 75 nm. This slice was rotated and projected using the same angle range and increment as for the real tomograms, resulting in a series of tilt images. These images were normalized, and Gaussian noise was added to the tilt images with signal-to-noise ratios of 1, 0.5,0.25, and 0.1. The tilt series were saved as MRC files, aligned, reconstructed, and denoised with the Warp model as was done for the real tomograms (see *Tilt series acquisition, reconstruction, and denoising*).

### Influence of cutting on linker orientation and chromatin folding path

The orientation of tomographic reconstructions of sections with respect to the cutting direction was determined from the orientation of the knife marks (Al-Amoudi, Studer et al., 2005). Two co-initial vectors were defined on the surface of the section containing knife marks: vector C is parallel to the cutting direction and vector Nc is normal to the cutting direction. Then we defined the vector Ns that is normal to the section surface, and accordingly normal to C and Nc (see Supplementary Figure S6A).

The end-to-end vector of each linker was projected onto C, N_c_ and N_s_ The projection values were used to test the null hypothesis *H*_0_: “the projection follows a uniform distribution”. As the projection value of isotropic random vectors onto any direction is uniform, this test is equivalent to testing the isotropy of the distribution of the linker end-to-end vectors. Generating isotropic random vectors is equivalent to drawing random points on the unit sphere (https://mathworld.wolfram.com/SpherePointPicking.html).

## DATA AVAILABILITY

Cryo-EM density maps for subtomogram averages of nucleosomes were deposited in the Electron Microscopy Data Bank (EMD-15480, EMD-15481, EMD-15483). The raw and denoised subvolmes of tomographic reconstructions of cHC and ECfHC were deposited in the Electron Microscopy Public Image Archive (EMPIAR-XXXX) and released upon publication.

## AUTHOR CONTRIBUTIONS

Conceptualization: ME, AL, FF

Methodology: ME, AL, JMV, FF, PC, DG, FT, WH

Investigation: FF, PC, DG, FT, ME, AL, JMV

Formal analysis: PC, JMV

Funding acquisition: ME, AL, JB

Supervision: ME

Writing – original draft: ME, AL, FF, JMV, PC

Writing – review & editing: ME, AL, FF, FT, JMV, PC, JB

## ACKNOWLEDGEMENTS

We are grateful to J. Dubochet for inspiring discussions and P.Schultz for critical reading of the manuscript. We thank H. Gnaegi (Diatome) for providing diamond knives for vitreous cryo-sectioning, Mohamad Harastani and Slavica Jonic (IMPMC) for support with subtomogram averaging. ME and DG thank A. Frangakis for providing the laboratory space and access to the equipment. We thank I.Tanoz for simulation of EM densities. FF and ME were supported by Centre for Integrative Biology (CBI), CNRS, Inserm, University of Strasbourg. FF, FT, AL and ME were supported by the French Infrastructure for Integrated Structural Biology (FRISBI ANR-10-INSB-005) and Instruct-ERIC. FT and AL were supported by iNext-Discovery, grant number 871037, funded by the Horizon 2020 program of the European Commission. This work benefited from discussions and workshops of the GDR ADN&G (CNRS GDR3536).

## FUNDING

French National Research Agency (ANR-20-CE11-0020-01 to AL, ANR-20-CE11-0020-02 to ME, ANR-20-CE11-0020-04 to JMV)

German Research Foundation (DFG EL 861/1 to ME)

German Federal Ministry of Education and Research (BMBF 02NUK054A to BJ)

